# Mitochondria Clearance Enables Macrophage-Driven Maturation of iPSC-Derived Cardiomyocyte Metabolism

**DOI:** 10.1101/2025.09.02.673264

**Authors:** Frank Ketchum, Lara Celebi, Lauren Hawthorne, Pinar Zorlutuna

## Abstract

Generation of functional engineered myocardial tissue remains a challenge, owing in part to lacking maturity of stem cell-derived cardiomyocytes. Current strategies to mature these cells fall short of achieving in vivo-like physiology. Macrophages, members of the innate immune system, reside in the heart and exert positive effects on cardiomyocyte function. We hypothesized that developmentally informed addition of macrophages to cardiomyocytes would improve their maturity. While some recent studies have added macrophages to stem cell-derived models of the human myocardium, these previous approaches do not replicate the early colonization of the heart. Addition of macrophages to developing cardiomyocytes 8 days after the start of differentiation significantly alters cardiomyocyte behavior. We show that macrophages drive improvements in metabolic capabilities of cardiomyocytes. Developing cardiomyocytes shed lowly polarized mitochondria, adopt a new mitochondria network architecture, and develop more active mitophagy programs after >20 days coculture with macrophages. This interaction is dependent on macrophage MerTK reception of cardiomyocyte-derived mitochondria material. These results inform our understanding of the responsibility of macrophages in the development of the myocardium, and we hope that these interactions can be leveraged to produce more physiologically relevant models of the human myocardium.

## Introduction

While cardiovascular diseases remain the leading cause of death in the US and worldwide,^1^ tools to accurately model the myocardium and recapitulate disease lag behind clinical need. Electrical,^2,3^ mechanical,^3^ and metabolic^4^ stimuli have been applied to generate mature cardiomyocytes (CMs) from stem cell sources with robust contraction and ability to model disease, but these approaches fail to replicate the complexity of the development of CMs in vitro, including the immunological niche of the heart.

Macrophages (Mϕ) reside in the heart, where they exert effects on heart remodeling^5–7^ and regulate CM metabolism.^8^ We hypothesized that mimicking macrophage (Mϕ) colonization of the myocardium in development would aid the maturation of iPSC-derived CMs (iCMs).

The cardiomyocyte is responsible for the contraction of the heart, which beats roughly once every second – 86,400 times a day. To power a constant heartbeat, the heart is estimated to consume 6kg of ATP per day.^9^ ATP, the cell’s energy currency, is produced largely in the mitochondria. In CMs, most ATP is generated from oxidation of fatty acids (60-90%), with significant contribution from oxidative metabolism of glucose (10-30%), and very little from glycolysis (<5%).^10^ To produce enough ATP for the high energetic demand of the heart, roughly one third of CM volume is occupied by mitochondria.^11^

Mitophagy, the selective trafficking of mitochondria to the lysosome for recycling, plays a vital role in CM development and maintenance. In the first 3 weeks post birth (P1-P21), CMs in the murine myocardium undergo an intensive mitophagy program to recycle embryonic mitochondria, which are inefficient in oxidative phosphorylation, and generate new mitochondria, which are competent in fatty acid metabolism.^12^ This drives the shift in myocardial metabolism from the predominantly glycolytic mode of embryonic development towards the primarily fatty acid oxidation-reliant metabolism of the neonatal and adult heart. The adult heart is also highly active in mitophagy, with mitochondria being overturned roughly every 2 weeks.^13^ New mitochondria are simultaneously generated to compensate for the recycling of old mitochondria. PGC-1α is regarded as the master controller of mitochondrial biogenesis, being activated in response to increased metabolic demand.^14^ Indeed, stem cell-derived CMs generate new mitochondria in a PGC-1α-dependent manner.^15^

CMs feature highly organized structure, with extensive mitochondria networks, well-aligned sarcomeres, and organized cell-cell junctions. While they show promise as a tool to model myocardial physiology, iCMs fail to fully replicate these features of mature cells. Previous approaches have attempted to address the lacking maturity of iCMs. Supplying mechanical structure and electrical pacing to iCMs, replicating these characteristics of the heart as an organ, has been explored to “exercise” cells and coax them to mature. These approaches lead to improvements in CM gene expression, calcium handling, and force generation.^3^ Further approaches have applied chemical methods to encourage the development of an adult-like metabolism, which improved CM function and enabled expression of pathology that was not possible without metabolic maturation.^16^ Mitophagy has also been explored as an avenue for maturation of CMs, with dexamethasone increasing measures of CM maturation without distinct structural changes.^17^ These methods have generated cells with improved functional characteristics, but there remain shortcomings in generation of in vivo-like cells. In fact, none of these approaches address that the heart and cells within it develop with cues from other tissues in the body, notably the immune system.

Macrophages (Mϕ), phagocytes and members of the innate immune system, have been of recent interest due to their residence in many tissues, including the myocardium. Mϕ migrate into the heart as early as embryonic day E10.5^18^ and remain throughout life.^19,20^ Previous studies have reported that these early embryonic Mϕ direct myocardial angiogenesis^21^ and contribute to limited regeneration potential in neonates,^22^ but the developmental role of this colonization in embryonic and early development remains unclear. Later in life, Mϕ are indispensable in the regulation of CM metabolism, with depletion of cardiac Mϕ leading to accumulation of dysfunctional mitochondria and impairing myocardial function in mouse models.^8^ Mϕ surface receptors MerTK and TIM-4 are implicated in efferocytosis,^23^ the clearing of debris including mitochondria-derived vesicles. This clearance of debris is important to the healthy development of the myocardium: genetic elimination of MerTK causes decreased expression of Myh6,^7^ signifying impaired cardiomyocyte maturity.

Conventionally, tissue engineering has relied on sourcing individual cell types from pure populations, combining them only to generate the final tissue construct. Previous tissue engineered models have integrated macrophages into cardiac tissues using this approach.^24,25^ To this end, Mϕ integration into human myocardium models has demonstrated improved maturity and functionality, increasing sarcomere alignment, beating strength, and closely aligning gene expression profiles with that of the native human myocardium. While this approach to tissue engineering allows for fine quality control of component cells, this does not reflect how tissues develop in vivo, where cells, tissues, and organ systems communicate with one another throughout development. Given the importance of Mϕ in the metabolism of the adult heart and their presence in the developing heart, we hypothesized that adding Mϕ to developing iCM cultures would improve the metabolism of these cells through direct interactions with iCM mitochondria. Through these changes, we expected to improve functional metrics of iCM maturity and elucidate the responsibilities of cardiac Mϕ in the developing heart.

## Results

### Developmental Addition of Macrophages to iCMs Improves iCM Metabolism

To replicate colonization of the myocardium by macrophages in development, we added Mϕ to developing iCMs and conducted metabolic and structural analyses. Isogenic Mϕ and CMs – those from the same iPSC line – were generated from established protocols. iCMs were induced via inhibition of GSK-3, followed by Wnt inhibition (GiWi), and underwent metabolic maturation as described by Feyen et al.^4^ Embryonic-like Mϕ were generated from embryoid body-derived hemogenic endothelial cells.^26,27^ During our iCM maturation protocol, 3 steps were identified as critical steps that may induce mitophagy due to metabolic stress. On day 9, cells are cultured in glucose-free medium containing fatty acids. Similarly, day 17 introduces CMs to a glucose-free media to prepare for administration of lipid-rich metabolic maturation media, which the cells are cultured in beginning on day 20. We hypothesized that addition of embryonic-like Mϕ at each of these steps would result in significant changes in respiratory measures due to the metabolic stress put on the CMs and metabolic interactions between Mϕ and CMs. A schematic showing the differentiation and Mϕ additions is shown in 1.A.

After differentiation day 30, with addition of Mϕ at prescribed times (fig. 1.A), iCMs were replated and Seahorse Mito Stress analysis was conducted. The oxygen consumption rate (OCR) values measured over time are shown in 1.B, with the coculture beginning at iCM differentiation day 9 showing increased OCR values. Processing of the OCR data indicated significantly increased basal respiration, maximal respiration, and spare respiratory capacity in iCMs cocultured with macrophages beginning on differentiation day 9. These differences were not observed in cultures where macrophages were added on days 17 or 20 (1.D). With an increase in metabolic activity, respiratory dysfunction was a concern. We checked for changes in proton leak, which correlates with increased reactive oxygen species (ROS) generation,^28^ and did not observe significant changes.

**Figure 1.**
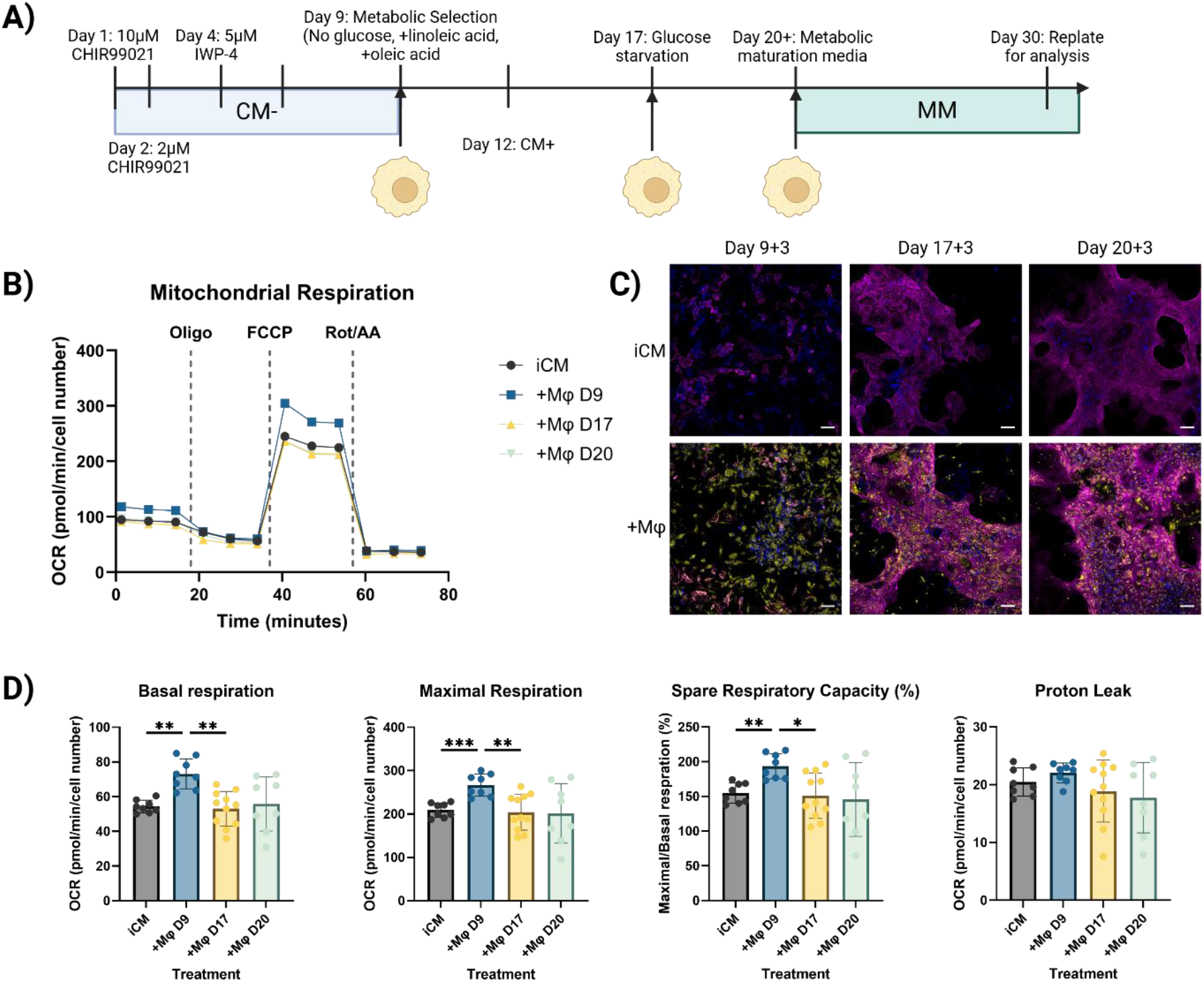
Developmentally informed addition of macrophages to iPSC-derived cardiomyocytes improves iCM metabolism. A) Schematic of macrophage addition to iCM differentiation and maturation protocol. B) Representative Oxygen Consumption Rate (OCR) plot from a mito stress test. C) Cepresentative immunofluorescence images of cocultures 3 days after macrophage addition. Yellow: CD14 (myeloid marker), Magenta: sarcomeric alpha actinin (cardiomyocyte), Blue: DAPI (nucleus). Scale bar: 50um. D) Basal and maximal OCR rates, spare respiratory capacity, and proton leak of day 30 iCMs with macrophage addition at days 9, 17, or 20 or no coculture. Mean ± SD. *: p<0.05, **: p<0.01, ***: p<0.001, Tukey’s t test with Dunnett’s correction.

To ensure that macrophages were integrating into and interacting with differentiating CM clusters, samples were fixed and stained 3 days after each addition of Mϕ, at days 12, 20, and 23. Mϕ clearly persist in the culture and associate with CMs, as indicated by CD14 presence in regions positive for sarcomeric α-actinin (1.C).

### Early Macrophages Modify iCM Mitochondria Network Architecture

Addition of macrophages to iCMs at differentiation day 9 had significant effects on the ability of iCMs to produce ATP. Because of this change, we selected day 9 as the critical time to add Mϕ to iCMs, and we investigated the changes in the mitochondria of these cells. Day 30 iCM cultures with or without the addition of Mϕ at day 9 were fixed and stained for Sarcomeric α-actinin, a component of the cardiac sarcomere, and COX4, a component of the electron transport chain, to visualize mitochondria network and sarcomere structure (2.A). Images of mitochondria in actinin-positive cells were analyzed using the publicly available MiNA pipeline^29^. While the total network size per image remained largely similar between CMs with and without coculture with Mϕ (2.B), measures of the degree of mitochondria network branching were significantly decreased by the addition of Mϕ early in CM development. Mean branch length, a measure of mitochondrion size, was significantly reduced in cocultured iCMs. Similarly, the mean total network length was reduced (2.C). This indicates a shift from more fused mitochondria to more punctate, dispersed mitochondria. Fission is a vital process in the dynamics of mitochondria and allows for the induction of mitophagy. The number of branches per network was significantly decreased in this condition (2.D), further supporting the observation of less extensive and interconnected mitochondria networks in the cocultured iCMs. Z-band spacing of each sarcomere was also measured from these images. In the human heart, sarcomeres lengthen from 1.7 to 2.2 microns as they mature.^30^ Our iCMs without Mϕ addition had a mean sarcomere length of 1.7µm, which falls just on the lower end of this range. CMs cocultured with Mϕ had an average sarcomere length of 2µm, approaching the length of a mature sarcomere (2.E).

### Cocultured iCMs Retain Highly Polarized Mitochondria

To further interrogate mitochondria function in iCMs with coculture with Mϕ, the voltage-sensitive dye JC-1 was employed. JC-1 changes emission based on mitochondrial polarization, with more red fluorescence signifying the presence of more highly polarized and therefore more active mitochondria. The ratio of red to green fluorescence intensity indicates the average membrane potential of mitochondria in the culture. We added Mϕ to mature iCMs to investigate changes in mitochondrial polarization. The JC-1 ratio significantly increased at 9 and 12 days in coculture versus iCMs alone (2.F). In mature iCMs cocultured with Mϕ, green JC-1 fluorescence, a measure of total mitochondria content, significantly decreased 9 days after addition of Mϕ, while the red signal did not change with coculture (2.G). This is apparent in the representative images (2.H). This indicates that while total mitochondria content in cocultured iCMs decreases acutely, highly polarized mitochondria are retained.

### Cocultured iCMs have Normalized Calcium Handling

To determine changes in functioning of the cocultured iCMs, we visualized calcium transients in mature iCMs after coculture with iMϕ. Action potential duration was unchanged by Mϕ coculture (2.I), but we observed significantly decreased variability of interpeak spacing (2.J), suggesting Mϕ regularized iCM calcium handling.

Taken together with the results of the Mito Stress Test, these results show that in the presence of Mϕ, CMs regulate their mitochondria networks to be more disperse and retain respiratory competent mitochondria while showing signs of elevated maturity.

### Macrophages collect and destroy iCM mitochondria

After addition of Mϕ to developing iCMs showed significant changes in mitochondria structure and function, we sought to assess the potential of human cardiac macrophages for pruning of iCM mitochondria as seen in mice^8^. To investigate direct transfer of mitochondria from iCMs to Mϕ, mitochondria originating from iCMs were stained with MitoTracker Red. To these iCMs with tagged mitochondria, iMϕ stained with CellTracker Green were added. These Mϕ contacted and engulfed CM-derived mitochondria within 48 hours (3.A). We also generated iCMs from a mtKeima-expressing human iPSC cell line. mtKeima is a mitochondria-targeted fluorescent protein. Here, mtKeima-expressing CMs and nonfluorescent Mϕ were cocultured, and we saw iCM-originated mitochondria passed to Mϕ (2.B and supplemental video).

### Macrophages in the Human Heart Overturn Myocardial Mitochondria

To verify that this mitochondria collection behavior is exhibited in human cardiac Mϕ, we collected human heart tissue slices. We stained these slices for CD68, a cardiac Mϕ marker, LAMP1, a lysosome marker, and COX4, a protein resident to the mitochondrial matrix (3.C). Imaging these tissue slices showed appreciable localization of COX4 to CD68-positive lysosomes. We calculate 4.9% (±3.6%) of COX4 fluorescence localized to CD68+/LAMP1+ lysosomes in these images, indicating a high turnover of mitochondria in the human myocardium by cardiac Mϕ. There was a distinct lack of LAMP1-positive regions that lacked CD68 expression.

### Gene Expression Changes are Not Responsible for Acute Changes in Mitochondria Networks

Because we hypothesize mitophagy to be a main driver of the maturation of iCMs in these cocultures, consistent with previous experiments, we interrogated expression of genes related to mitophagy and mitochondria biogenesis over the length of CM maturation. Parkin (*park2*) is an adaptor protein responsible for initiation of the phagophore which engulfs a mitochondrion to initiate mitophagy,^31^ and DRP1 (*dnm1l*) drives mitochondria fission. Mitochondria biogenesis was measured through the gene *atp5b*, a nuclear-encoded subunit of the F^1^ region of ATP synthase. Interestingly, *park2, dlp1*, and *atp5b* all tracked well with one another over the length of the differentiation regardless of the addition of macrophages to the culture. At differentiation day 30, however, parkin, dlp1, and ATP synthase expression were each significantly upregulated in the coculture condition (4.A), indicating increased mitochondria turnover through mitophagy and replenishment by de novo synthesis.

### mtKeima Reveals Changes in Mitophagy Flux due to Mϕ Coculture

Distinct changes in mitochondria network structure shown in Figure 2 were reflected by changes in gene expression only at day 30 of the differentiation, 21 days after the addition of Mϕ. To determine mitophagic flux in these differentiating cells at a protein level, we employed an iPSC line expressing mtKeima. mtKeima includes a mitochondrion transport signal sequence, causing selective trafficking to the mitochondria. When trafficked to the lysosome during mitophagy, the low pH environment causes a shift in the excitation to a redder wavelength. This cell line has previously been employed to measure mitophagy in iCMs.^32^ iCM differentiations expressing mtKeima were cocultured with macrophages without this fluorescent tag to specifically interrogate the dynamics of iCM mitochondria. We measured mtKeima signal in iCMs over the course of the differentiation, with and without addition of Mϕ at day 9. Coculture with macrophages increased the ratio of red to green mtKeima signal, indicating an increase in mitophagic flux, at days 25 and 30 (fig 4C). Mitophagic flux was at its highest in both groups at differentiation day 12, at the end of the metabolic purification step.

**Figure 2.**
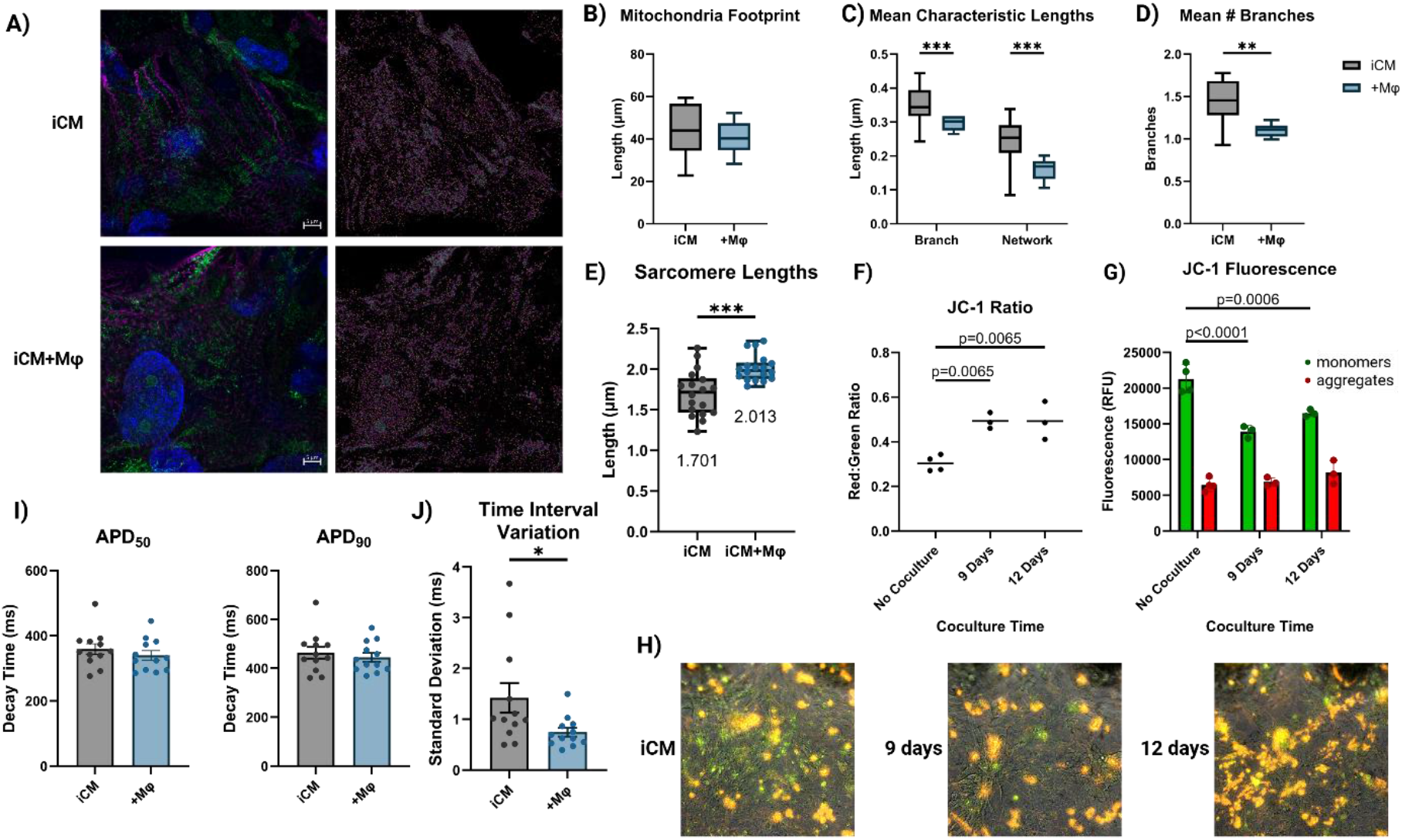
Addition of macrophages to iCMs in development causes maturation of subcellular structures. A) Representative images of day 30 iCMs, with or without addition of Mϕ at day 9, stained with COX4 (mitochondria, green), Sarcomeric alpha-actinin (CM, magenta), and DAPI (nucleus, blue) B) Quantification of mitochondria network footprint, C) mean branch and total network length, and D) mean number of branches per network in day 30 iCMs with or without addition of Mϕ at day 9. E) Measured sarcomere lengths in images. F) JC-1 ratio in mature iCMs cocultured with Mϕ for 9 or 12 days. G) Raw JC-1 fluorescence values in iCMs with or without coculture. H) Representative images of JC-1 staining showing changes in green and red fluorescence. Calcium transient duration (I) and peak-to-peak interval variation (J) in iCMs with or without addition of macrophages at day 30. *: p<0.05, **: p<0.01, ***: p<0.001, p values noted from Tukey’s t test.

**Figure 3.**
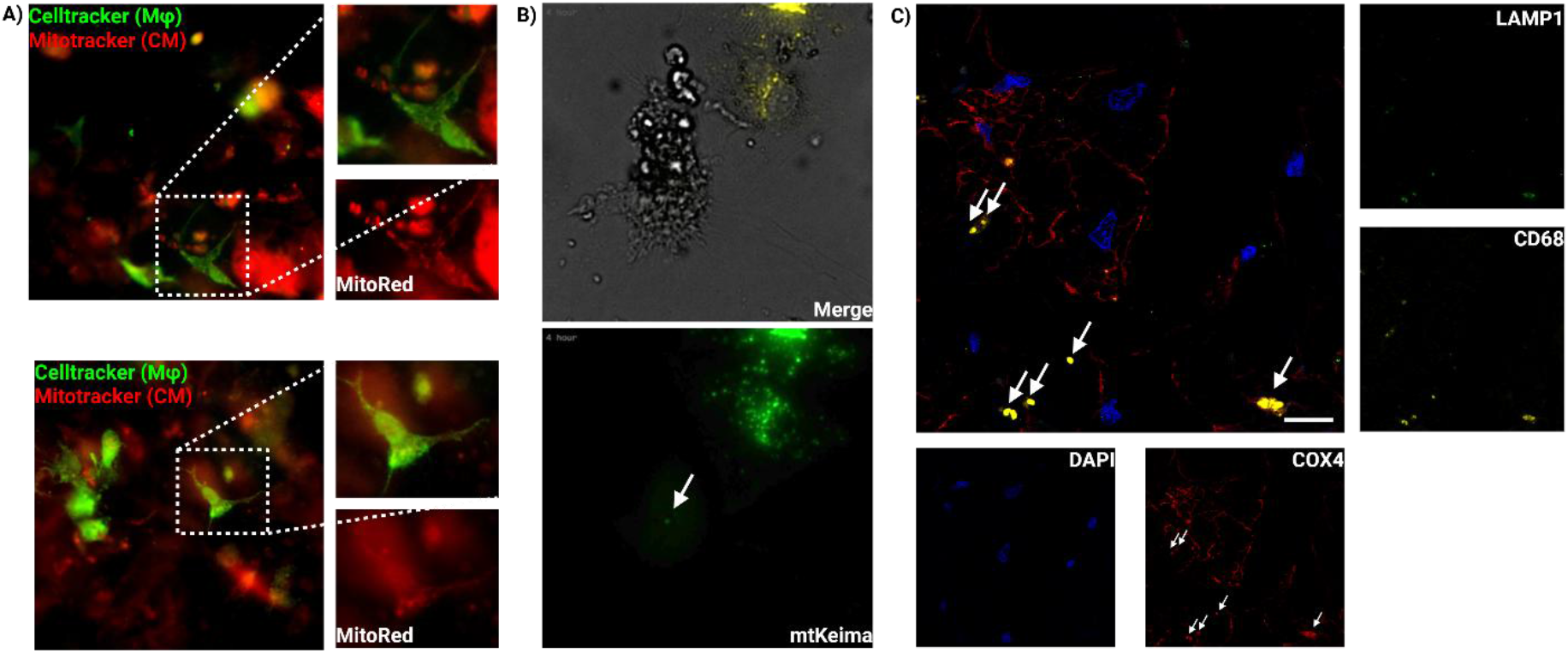
Macrophages collect and destroy iCM-derived mitochondria. A) Airyscan images showing uptake of mitochondria by macrophages in coculture. Red: MitoTracker Red (iCMs), green: CellTracker Green (iMϕ). B) Frame from timelapse image of iCM-iMϕ coculture showing transfer of mtKeima-positive mitochondrion from iCM to iMϕ. Arrow indicates mtKeima-positive iCM mitochondrion within Mϕ. Green: mtKeima (iCM). C) Immunohistochemistry images of human heart slices showing macrophages destroying mitochondria. Blue: DAPI (nucleus), yellow: CD68 (Mϕ), green: LAMP1 (lysosome), red: COX4 (mitochondria). Arrows indicate COX4 localized to CD68+/LAMP1+ Mϕ lysosomes. Scale bar: 10µm.

**Figure 4.**
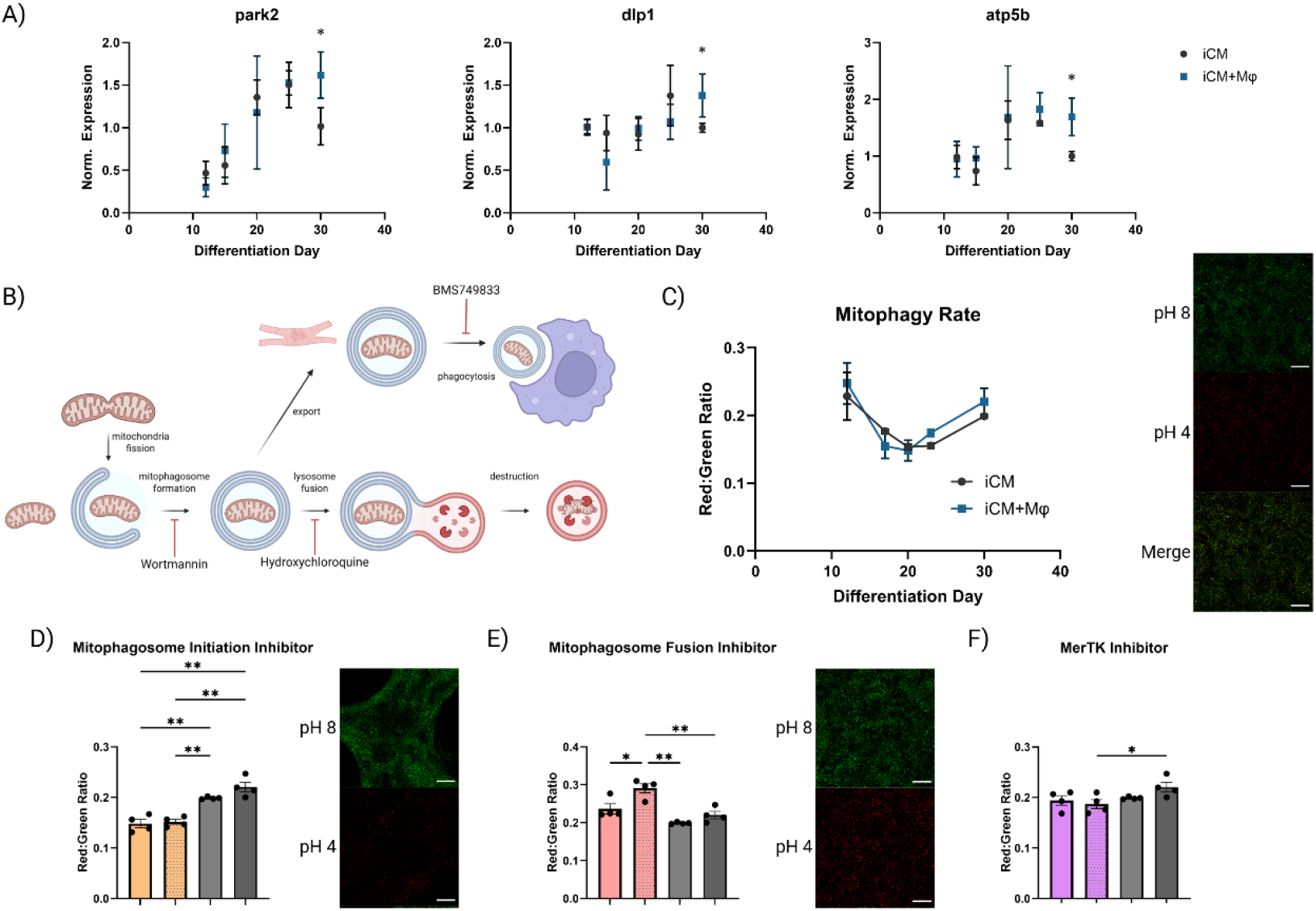
Mitophagy and mitochondria biogenesis are stimulated at different times in cocultures. A) Results of PCR from samples taken at days 12, 15, 20, 25, and 30 of iCM differentiation with or without macrophage addition at day 9. *: p<0.05, Tukey’s t test. B) General schematic of mitophagy steps with inhibitors. C) Mitophagy flux in mtKeima-expressing iCMs as measured by ratio of low-pH mitochondria (red) to normal pH mitochondria (green) at days 12, 17, 20, 23, and 30 of iCM differentiation with or without Mϕ addition at day 9. Right: representative images. Graphs of mitophagy flux of iCMs at day 30 with or without Mϕ addition at day 9 and with or without wortmannin treatment (D), hydroxychloroquine treatment (E), or BMS794833 treatment (F) beginning at day 9. Right: representative images. *: p<0.05, **: p<0.01, Tukey’s t test. Scale bar: 100 µm for all images.

**Figure 5.**
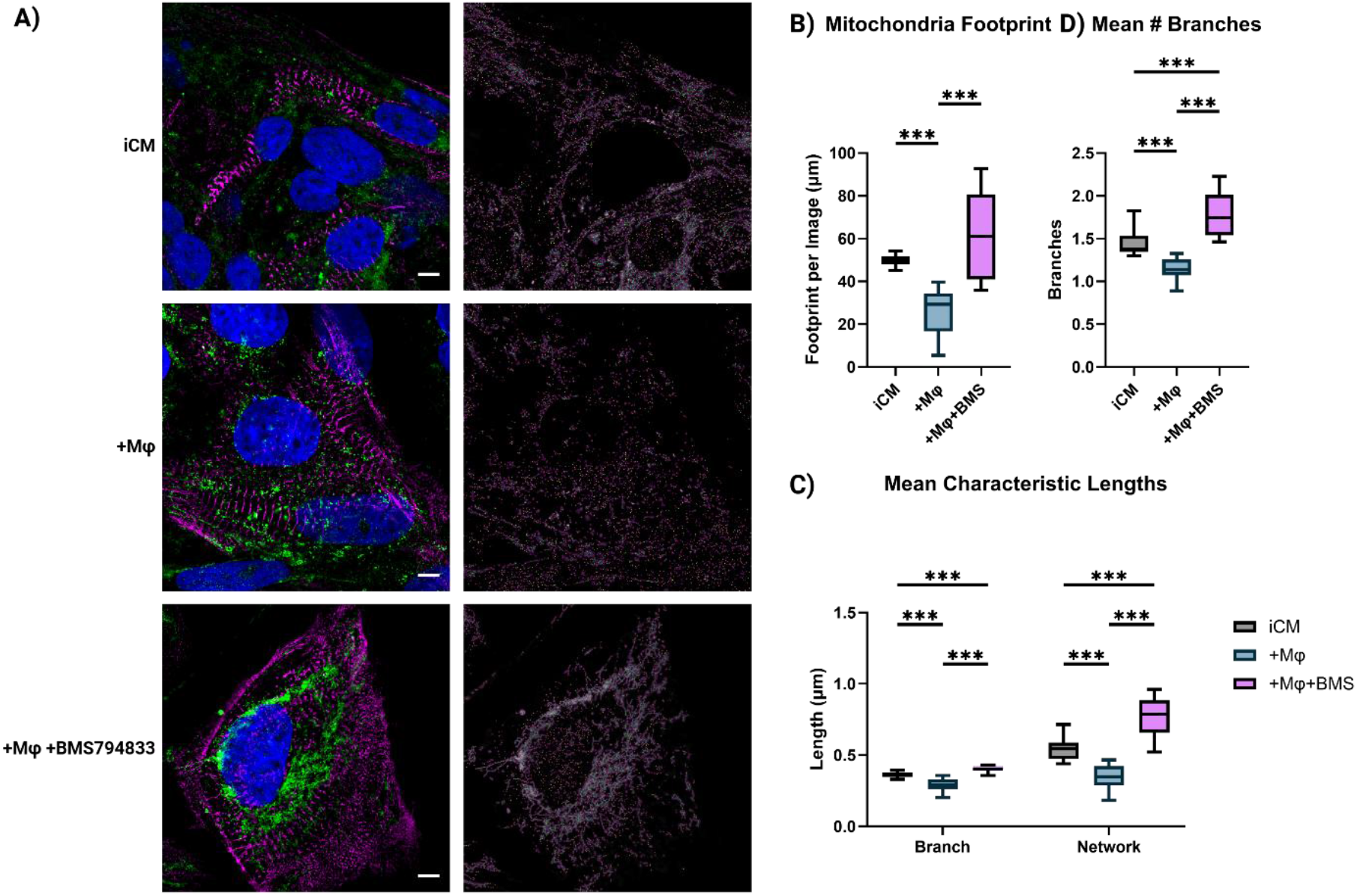
MerTK function is necessary for macrophage-driven maturation of iCMs. A) Representative images showing Z discs (Sarcomeric α-Actinin, magenta), mitochondria (COX4, green), and nuclei (DAPI, blue) (left), and mitochondria network analysis (right) of iCMs in each condition. Scale bar: 5µm. B) Mitochondria network metrics from immunofluorescence images of iCMs cocultured with iMϕ with or without inhibition of MerTK by BMS749833. *: p<0.05, **: p<0.01, ***: p<0.001, Tukey’s t test with Dunnett’s correction.

### Inhibition of Mitophagy and Mitochondria Uptake Prevents Mϕ Effects on iCM Mitophagy

We used this mtKeima system to determine the mechanism of Mϕ effects on iCM mitophagy. To iCM cultures with and without Mϕ addition, we added drugs previously shown to influence iCM mitophagy. Wortmannin disrupts the formation of the autophagosome,^33^ hydroxychloroquine disrupts the fusion of the autophagosome with the lysosome,^33^ and BMS794833 inhibits MerTK,^34^ a scavenger receptor on Mϕ which interacts with mitochondria-derived vesicles (MDVs).^7^ Figure 4.B illustrates the function of these drugs and how they interact with mitophagy and mitophagosome uptake by macrophages. Wortmannin significantly decreased the mitophagy flux in iCMs regardless of the addition of Mϕ, as expected (4.D). Hydroxychloroquine, which alkalizes the lysosome and prevents its function, increased the proportion of low-pH mitochondria (4.E), likely due to the accumulation of low-pH autophagosomes containing mitochondria in both iCMs and Mϕ.^35^ This effect was exacerbated by coculture with Mϕ, indicating that Mϕ increase mitophagy flux and completion of mitophagy with the degradation of mitochondria is not required for Mϕ to affect iCM mitophagy. Interestingly, the addition of BMS794833, an inhibitor of the scavenger receptor MerTK, completely abrogated macrophages’ effect on iCM mitophagy (4.F). The treatment with this MerTK inhibitor and Mϕ led to a significantly lower mitophagic flux than Mϕ addition alone. MerTK inhibition had no effect on iCMs alone. Because increased mitophagy was not observed with Mϕ addition in groups treated with wortmannin or BMS734899, we argue that Mϕ uptake of mitochondria through MerTK is required for iMϕ to stimulate iCM mitophagy.

### MerTK Function is Required for iCM Mitochondria Network Remodeling

Based on the abrogation of enhanced mitophagy in iCMs treated with iMϕ, we sought to investigate MerTK’s importance in the interaction between these cells. iCMs were differentiated and macrophages were added on D9, with or without the addition of the MerTK inhibitor BMS749833. Cells treated with BMS794833 were treated for the entire duration of the differentiation (starting on D9 when the macrophages were added). Cells were fixed and stained after day 30 as before. Measurement of the mitochondria networks in these iCMs showed that Inhibition of MerTK in cocultures resulted in a further increase of mitochondria network size and connectivity. Each of the pairwise comparisons showed statistical significance. This shows that macrophages promote mitochondria biogenesis in iCMs without the need for MerTK interactions or mitochondria scavenging, while MerTK is essential for the phagocytic removal of mitophagosomes and promotion of mitophagy. This is consistent with previous results showing decreased mitophagy in cocultured iCMs with the inhibition of MerTK.

## Discussion

Our results demonstrate that early coculture with macrophages directly improves iPSC-derived cardiomyocyte (iCM) metabolic and structural maturity through MerTK-mediated scavenging of iCM mitochondria. This mimics macrophage colonization and mitophagy-driven maturation seen in perinatal development. Inhibition of MerTK abrogated this effect, illustrating the critical role of macrophage pruning of iCM mitochondria in this interaction.

Because Mϕ infiltrate the heart during development,^20,36^ reside throughout life,^19,20,37^ and regulate CM metabolism,^8^ we sought to leverage interactions between Mϕ and developing CMs to generate more mature CMs from iPSC sources. Consistent with our hypothesis, the addition of Mϕ to differentiation day 9 iCMs drastically increased mitochondrial respiration capacity after day 30, more than 20 days after Mϕ were introduced to these cells. The addition of Mϕ on differentiation day 9 also caused persistent changes in these cells consistent with increased maturity, with improved calcium handling, namely more regular calcium transients, and lengthened sarcomeres on day 30. Coinciding with this improved maturity, we observe distinctly different mitochondrial network geometry in CMs after adding Mϕ at day 9. Interestingly, other groups report that Mϕ addition to engineered tissues increases the maturity of these tissues and the CMs within,^24,25^ but these studies do not report increases in mitophagy-related genes despite the apparent importance of Mϕ in stimulating CM mitophagy.^8^ We argue that a more developmentally relevant addition of Mϕ to CMs better replicates the positive effects they exert on cardiac tissue in vitro.

The increase in CM metabolism with developmental Mϕ coculture coincides with a decrease in mitochondria network size and connectivity in the same conditions. The tubularity/interconnectedness of mitochondria networks serve to prevent mitophagy, whereas more fragmented networks are more active in mitophagy.^38^ Because of the importance of mitophagic clearance in CM development and maintenance, we believe that the cocultured CMs take on this altered network structure to allow the maturation and upkeep of their respiratory machinery. In support of this, the adult human heart is reported to contain more “fizzed” mitochondria which localize near sarcomeres.^12^ Taken together, iCMs cocultured with Mϕ beginning early in the differentiation take on a more structurally mature phenotype. The geometry of these networks allows for acute regulation of mitophagic flux, which is high in the myocardium, as well as energy generation and delivery to sarcomeres, where it is most demanded.

Expression of mitophagy gene *park2* indicates that on an expression level, mitophagy machinery is not upregulated by macrophages acutely, but rather much later (∼21 days). This is reflected in other reports’ lack of significant changes in gene expression of mitophagy-related genes with the incorporation of Mϕ.^24,25^ While this is the case, we did detect differences in mitophagy flux using the mtKeima reporter, with Mϕ significantly increasing mitophagic flux in CMs as early as 12 days after macrophage addition. We also observe a distinct change in mitochondria structure of iCMs by differentiation day 30 after addition of Mϕ at day 9. Given the long delay between Mϕ addition and gene expression changes, it appears that mitophagy is not modulated at a gene expression level acutely, but rather at a protein level, with gene expression increasing later to enable the continued regulation of modified mitochondria networks.

The exact mechanism behind these changes in CMs remains unclear. On a gene expression level, no acute changes in mitophagy (*park2, dlp1*) nor mitochondria biogenesis (*atp5b*) genes are responsible for changes in respiration later in the lives of these cells. The delay until detectable gene expression changes suggests that Mϕ stimulate reprogramming of CMs which causes them to be more active in mitophagy. Epigenetic changes may be responsible, or the altered mitochondria morphology we observe may perpetuate by self-stimulating mitophagy within CMs.

Macrophage scavenging of exported mitochondria-derived material is indispensable in the interaction between CMs and Mϕ. MerTK, a receptor tyrosine kinase involved in efferocytosis,^7^ is required for this process. In coculture, we see Mϕ take up CM-derived mitochondria, and human heart slices show an accumulation of mitochondria material in Mϕ lysosomes, which shows that this mechanism occurs in the human myocardium and is not limited to mice.

Pharmacological inhibition of MerTK in cocultures abrogated Mϕ-driven increases of mitophagy in CMs, returning flux to its baseline rate. Wortmannin, a broad inhibitor of autophagy,^33^ suppressed CM mitophagy and prevented Mϕ-mediated rescue of CM mitophagic flux. In both cases, Mϕ were rendered unable to take up CM-derived MDVs and failed to stimulate mitophagy in CMs.

Communication between CMs and Mϕ appears to be bidirectional. MerTK-reliant uptake of MDVs causes maturation of cardiac Mϕ, which then stimulate CMs to increase mitophagy. MerTK signaling stimulates M2 polarization of Mϕ,^39^ whose effects on angiogenesis and tissue remodeling resemble cardiac Mϕ phenotype.

Further supporting this mechanism, Mϕ increased CM mitophagy flux in hydroxychloroquine (HCQ) treated cultures. HCQ alkalizes the lysosome, preventing autophagosome degradation.^33^ HCQ increased the proportion of low-pH mitochondria in CMs, which we interpret as an accumulation of mitolysosomes in these cells. With Mϕ coculture, this effect was exacerbated; Mϕ still promoted increased mitophagy in CMs. This contrasts with MerTK and wortmannin inhibited cultures, where Mϕ were unable to stimulate mitophagy in CMs. We theorize that intact MerTK signaling in the HCQ case allowed Mϕ to stimulate increased mitophagic flux in CMs despite inability to digest MDVs.

Through mitochondria structure analysis, we see an accumulation of mitochondria in CMs when cocultured with Mϕ while MerTK is inhibited. We did observe increased ATP synthase gene expression in cocultured iCMs, showing that Mϕ promote mitochondrial biogenesis in CMs. This agrees with previous studies.^24,25^ The accumulation of mitochondria in the MerTK-inhibited cocultures supports this and shows that the stimulation of mitochondria biogenesis and mitophagy by Mϕ are regulated through different mechanisms, where increased mitochondria biogenesis is not a response to increased mitophagy, but stimulated independently by Mϕ. Interestingly, when MerTK was inhibited, CMs contained more mitochondria, similar to mouse hearts with depleted Mϕ.^8^ The observed decrease in mitophagy flux with MerTK inhibition is likely the cause of mitochondria retention.

Research applying animal models lacking embryonic cardiac macrophages to heart development is limited. One study has determined that *op/op* mice, those lacking functional M-CSF, have impaired myocardial angiogenesis and coronary artery development.^21^ In fact, these mice don’t completely lack macrophages, confounding findings.^40^ Through our studies, we elucidate some effects of Mϕ on human CM development. We observe Mϕ directly interacting with CM mitophagy, which is indispensable in the development of a mature heart metabolism.^12^ In developing CMs, mitophagic flux peaks by differentiation day 12, when CMs are cultured in media with fatty acids and without glucose. Addition of Mϕ to CMs before this peak lead to increased metabolic maturity in CMs. The decreased mitochondria presence in iCMs after coculture with Mϕ correlates with the heart’s metabolic switch; JC-1 results show lowly polarized embryonic mitochondria are recycled in iCMs when in culture with Mϕ. This supports our theory that Mϕ are essential not only for CMs to regulate their mitochondria networks in homeostasis but also in development for the overturning of embryonic, fatty acid metabolism-incompetent mitochondria and acquisition of mature metabolism. Further research in vivo may help establish these responsibilities of Mϕ in the developing heart once a suitable model system is developed.

Collectively, our results position Mϕ treatment as a developmentally informed strategy for enhancing iCM maturity. Beyond improving in vitro modeling, this approach may inform regenerative therapies that harness immune cues to support cardiac repair and remodeling.

## Conclusion

Here we show that the addition of macrophages to iPSC differentiation towards cardiomyocytes improves measures of CM respiration and maturity. MerTK-dependent clearance of CM mitochondria is required for these effects. This suggests a developmental role of macrophages in the stimulation of mitochondria network maturation in CMs. By leveraging this crosstalk, we move closer to developing methods for generation of mature CMs from iPSC sources.

## Methods

### Cell lines and iPSC maintenance

DiPS 1016 SeVA: male dermal fibroblast-derived iPSC

MTK-11: mtKeima-expressing iPSCs were generated via CRISPR-Cas9 gene integration as described previously^32^ from MS19-ES-H (female, PBMC-derived iPSC) and provided by Professor Nuo Sun.

iPSCs were maintained in mTeSR Plus media (StemCell) with media change at least every other day. Cells (>70% confluency) were passaged using accutase (StemCell) and plated on geltrex (Fisher) coated plates in mTeSR Plus with 5µM ROCK inhibitor Y-27632. Mycoplasma testing was performed quarterly.

### iCM differentiation

Differentiation of iCMs from iPSCs was achieved by a modified GiWi protocol. iPSCs (DiPS 1016 SeVA or MTK-11) at >90% confluence were cultured in CM-(RPMI 1640 (Corning) with 1x B27 without insulin (Gibco) + 2µM β-ME + 1% Penicillin) with 10µM CHIR99021 (StemCell) on day 1. 24 hours later and the following day, the media was changed to CM-with 2µM CHIR. On day 4, cells were cultured in CM-with 5µM IWP-4 (StemCell). 48 hours later, the media was changed to CM-. Metabolic maturation of iCMs was conducted according to Feyen et al^4^. On day 9, iCMs were purified through culture in RPMI 1640 without glucose with 1xB27 + 1x Linoleic acid-Oleic acid albumin (Sigma-Aldrich/Merck) for 3 days. CM+ (RPMI 1640 (Corning) with 1x B27 (Gibco) + 2µM β-ME + 1% Penicillin) was used for days 12-17. On day 17, iCMs were prepared for metabolic maturation through culture in RPMI 1640 without glucose with 1xB27 for 3 days, followed by culture in metabolic maturation media^4^ until analysis.

### iMϕ differentiation

Macrophages were generated from DiPS 1016 SeVA as described by van Wilgenburg et al.^27^ and Buchrieser et al.^26^ Briefly, mechanically dissociated iPSCs were differentiated in mTeSR plus media supplemented with VEGF, SCF, and BMP-4 for 4 days in low-attachment plates, followed by plating on Matrigel (Corning)-coated plates. Adherent EBs were then maintained in X-VIVO 15 + 50ng/ml M-CSF + 25ng/ml IL-3 for up to 3 months, with collection of suspended macrophage precursors every 3-4 days.

### Addition of iMϕ to CM differentiations

On days 9, 17, or 20, macrophages were freshly collected from EBs and rinsed with PBS. Macrophage precursors were resuspended in appropriate iCM media supplemented with 50ng/ml M-CSF and added to iCMs as a fresh media change.

### Seahorse mitochondria stress assay

On day 30, iCMs were replated in fibronectin (Sigma)-coated 96-well seahorse plates (Agilent) by harvesting with trypsin-EDTA. Pelleted iCMs were resuspended in CM+ with 10% KOSR (Thermo) and 5µM Y-27632 (StemCell). Replated cells were then cultured for 4 days in metabolic maturation media before analysis. One hour before the assay, the culture medium was replaced with Agilent Seahorse XF RPMI Basal Medium with 2 mM Glutamine, 10 mM glucose, and 1 mM sodium pyruvate. The mitochondrial stress assay was performed as prescribed by the manufacturer with sequential addition of oligomycin (2.5 µM final concentration), FCCP (2.0 µM final concentration), and rotenone (2.5 µM final concentration)/antimycin-A (2.5 µM final concentration) solutions. iCM number was quantified using Hoescht 33342 (Thermo) staining, followed by analysis in ImageJ. OCR was normalized to cell number. Basal respiration, maximal respiration, spare respiratory capacity, and proton leak were analyzed in Seahorse Analytics.

### Calcium Flux Analysis

Cells were washed with PBS, and the medium was replaced with a Ca^2+^ -sensitive Calbryte 520 AM following the manufacturer’s instructions. After 30 minutes of incubation, the medium was replaced with Tyrode’s solution. Real-time beating videos of iCMs were recorded using a fluorescence microscope equipped with a Hamamatsu C11440 digital camera, with a 100 ms exposure time for 20 seconds. Variability of interpeak spacing was quantified as the standard deviation of the interbeat intervals.

### Immunofluorescence staining

Cells on glass cover slides or tissue culture plastic or sectioned tissues were fixed in 4% PFA (Electron Microscopy Solutions) solution for 15 minutes. Fixed samples were permeablized with 0.1% Triton-X 100 in water for 15 minutes followed by three rinses in PBS and blocking in 5% goat serum for 30 minutes at room temperature. Primary antibodies were added directly to blocking solution and samples were incubated at 4C on a rocker overnight.

Secondary staining was performed by incubating samples in 1% BSA solution with 1:2000 secondary antibody at room temperature in the dark for 1 hour. Nuclear counterstaining was performed using DAPI in PBS for 10 minutes. Tagged primary antibodies (CD68-Alexa Fluor 594) were added after counterstaining. Glass coverslips were transferred to glass slides and mounted using Prolog Antifade Gold reagent (Thermo). Imaging was performed on a Zeiss LSM900 /AxioObserver 7 microscope with 20x objective in air or oil immersion 63x objective. Airyscan images were generated using an Airyscan 2 module.

### Mitochondria network analysis

Immunostained samples were imaged on glass using Airyscan on a 63x oil objective. The COX IV channel was isolated, and the MiNA pipeline^29^ was used to analyze the structure of the mitochondria networks. Parameters used in the analysis were as follows: unsharp mask: r=10, w=0.5; CLAHE: b=127, bins=256, max slope=3; median fill: r=2, as prescribed.

### Sarcomere Length Quantification

Immunostained samples were imaged on glass using Airyscan on a 63x oil objective. ImageJ was used to measure fluorescence signal in the Sarcomeric α-Actinin channel along the length of a sarcomere. A custom Python code was employed to measure the average peak-to-peak distance.

### JC-1 assay

Cells were plated in 96-well, black-walled culture plates coated with fibronectin. At the end of the experiment, cells were washed twice with PBS, and JC-1 (Thermo) solution (10µM in PBS) was added to the cells. Cells were incubated in JC-1 solution for 10 minutes at 37C in the dark.

### Human Samples

De-identified human hearts that were deemed unsuitable for transplantation and donated to research were acquired from the Indiana Donor Network; IRB approval was waived as the Indiana Donor Network did not provide any identifying information. Hearts from donors were cut into region-specific pieces and stored at −80 °C until further use.

### Polymerase Chain Reaction

RNA was collected from adherent iCM cultures using the Qiagen RNEasy Kit, per manufacturer’s protocol. RNA concentration and purity was quantified by NanoDrop 2000 spectrophotometer. RNA was stored at -80C until use.

RNA concentrations of samples were normalized before first strand cDNA synthesis by dilution in RNAse-free water. cDNA was synthesized using the SuperScript™ VILO™ cDNA Synthesis Kit (Thermo) per manufacturer’s protocol.

cDNA was amplified using TaqMan Fast Advanced Master Mix (Thermo) and respective primer according to manufacturer’s specification.

### Pharmacological inhibition of mitophagy and measurement

iCMs were generated from MTK-11 iPSCs, with or without addition of DiPS 1016 SeVA-derived macrophages on day 9, as described above. Beginning on differentiation day 9, culture media was supplemented with mitophagy drugs. Concentrations used are shown in **Table 2**. Mitophagy-modulating drugs employed in experiments.**Table 2**.

**Table 1.**
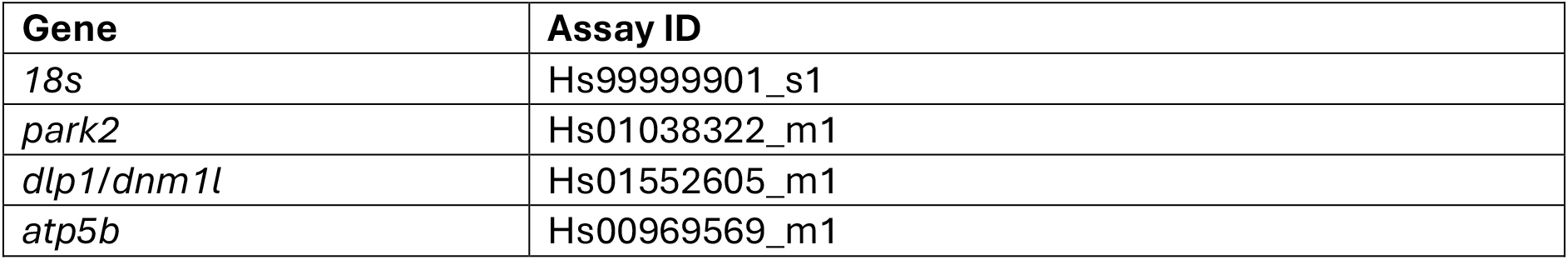
PCR Primers employed in experiments.

| Gene | Assay ID |
| --- | --- |
| *18s* | Hs99999901_s1 |
| *park2* | Hs01038322_m1 |
| *dlp1/dnm1l* | Hs01552605_m1 |
| *atp5b* | Hs00969569_m1 |

**Table 2.** Mitophagy-modulating drugs employed in experiments.

| Drug | Working Concentration | Vehicle |
| --- | --- | --- |
| Wortmannin | 0.2 $\mu\text{M}$<sup>41</sup> | DMSO |
| Hydroxychloroquine | 20 $\mu\text{M}$<sup>42</sup> | Water |
| BMS794833 | 1 $\mu\text{M}$<sup>34</sup> | DMSO |

At prescribed times, media was removed from wells and cells were washed once with PBS, then fresh PBS was added. 20x confocal images were taken of each well, and fresh media containing appropriate drugs was added to cells. Mitophagy flux was quantified by the ratio of fluorescence intensity stimulated by 488nm laser divided by the fluorescence intensity stimulated by 594nm laser.

### Statistics and Reproducibility

All statistical testing was conducted in PRISM. Statistical tests used are noted where applicable.

## Acknowledgements

We would like to thank Nuo Sun from the Department of Physiology and Cell Biology at The Ohio State University Wexner Medical Center for providing the mtKeima-expressing iPSCs for live mitophagy measurements. We acknowledge the Notre Dame Optical Microscopy Core for providing the microscope used for tissue slice imaging. We acknowledge the Biological Screening and Development Core for providing the Seahorse XFe analyzer for metabolic activity measurements. This work was funded by NSF EFRI BEGIN OI Award number 2422333 and the Naughton Faculty Research Accelerator Grant. All figures were created in part in Biorender.

